# Genetic diversity and domestication of hazelnut (*Corylus avellana*) in Turkey

**DOI:** 10.1101/622027

**Authors:** Andrew J. Helmstetter, Nihal Oztolan-Erol, Stuart J. Lucas, Richard J. A. Buggs

## Abstract

- Assessing and describing genetic diversity in crop plants is a crucial first step towards their improvement. The European hazelnut, *Corylus avellana*, is one of the most economically important tree nut crops worldwide. It is primarily produced in Turkey where rural communities depend on it for their livelihoods. Despite this we know little about hazelnut’s domestication history and the genetic diversity it holds.
- We use double digest Restriction-site Associated DNA (ddRAD) sequencing to produce genome-wide dataset containing wild and domesticated hazelnut. We uncover patterns of population structure and diversity, determine levels of crop-wild gene flow and estimate the timing of key divergence events.
- We find that genetic clusters of cultivars do not reflect their given names and that there is limited evidence for a reduction in genetic diversity in domesticated individuals. Admixture has likely occurred multiple times between wild and domesticated hazelnut. Domesticates appear to have first diverged from their wild relatives during the Mesolithic.
- We provide the first genomic assessment of Turkish hazelnut diversity and suggest that it is currently in a partial stage of domestication. Our study provides a platform for further research that will protect this crop from the threats of climate change and an emerging fungal disease.

## INTRODUCTION

Understanding genetic diversity in crop plants and their wild relatives is critical for improving breeding programmes (Zamir, 2001), combatting disease (Zhu *et al*., 2000) and determining the impact of domestication (Wright, 2005). Advances in genomic sequencing and the generation of reference genomes have helped identify genetic variation associated with phenotypes important for agriculture (Bevan *et al*., 2017). Such approaches have been used to uncover the history and diversity of model crop species such as rice (He *et al*., 2011) and maize (van Heerwaarden *et al*., 2011). However, methods are available that can be used in non-model crop species to sequence across the entire genome cheaply and efficiently (Andrews *et al*., 2016). This has unlocked the potential for genomic studies in non-model crop species such as the Scarlett runner bean, *Phaseolus coccineus* (Guerra-García *et al*., 2017) and the curcurbit bottle gourd, *Lagenaria siceraria* (Xu *et al*., 2013). These approaches can be applied to crops that may not be widely cultivated but are critical to the economies and communities of developing regions. Improving our understanding of genetic diversity with genomic data can kick-start research towards crop improvement that will have a real and lasting impact on farmers and communities. One such economically important yet understudied crop is the European hazelnut, *Corylus avellana* L.

*Corylus avellana* is a hermaphroditic, self-incompatible shrub that is typically clonally propagated (Molnar, 2011). The nut of *C. avellana* is one of the most valuable tree nut crops worldwide yet we have relatively few resources relevant to its improvement as a crop species. Small proportions of the world’s hazelnut production comes from countries such as Spain, Azerbaijan and the USA while Italy produces approximately 15%. The vast majority, 70-80%, of the world’s hazelnut market is produced in Turkey (Gökirmak *et al*., 2008). It is Turkey’s largest agricultural export and 61% of the rural Black Sea population rely on smallholdings of hazelnut for their primary income (Gönenç et al., 2006), making the performance of the crop critical to the livelihood of the inhabitants of this region. However, spring frosts and summer droughts regularly reduce hazelnut yields by up to 85% (Ustaoglu, 2012) and this has knock-on effects on the local economy. Furthermore, a new powdery mildew disease has emerged in recent years, and is considered by Turkish producers to be the most significant immediate threat to hazelnut production. The disease is now recognized to be widespread across the eastern Black Sea region and 60-100% of trees have been found to be affected in areas close to sea level (Lucas et al., 2018). Despite the economic importance of this tree nut crop and the current threats it faces, we know little about genetic variation in wild and cultivated forms.

Previous studies have provided insight into diversity among cultivated and wild hazelnuts across Europe (e.g. (Boccacci *et al*., 2006; Gökirmak *et al*., 2008; Boccacci *et al*., 2013) as well as specifically in Turkey (Kafkas *et al*., 2009; Gürcan *et al*., 2010; Öztürk *et al*., 2017), using a small number of markers. Genome-wide studies have commenced on an American cultivated strain, primarily to understand resistance to the disease eastern Filbert blight (EFB) (Rowley *et al*., 2018). EFB is an important issue in the USA but additional work is needed where the crop is primarily produced if we are to maximize the social and economic impact of hazelnut research (Bacchetta *et al*., 2015).

In this study we aim to lay the groundwork for a genomic perspective on hazelnut in Turkey. We conduct double digest restriction-site associated DNA sequencing (Peterson *et al*., 2012) on more than 200 individuals, principally wild and cultivated *C. avellana* from the Black Sea region of Northern Turkey. To provide context in our genomic analyses we also include specimens from the UK, Georgia and the Campania region of Italy as well as samples from other members of the same genus, *C. colurna* and *C. maxima*. We use these genomic data to determine patterns of genetic diversity and structure among and within wild and cultivated populations.

Domestication is thought to cause a rapid reduction in population size, when early farmers isolate a strain, followed by expansion. This ‘domestication bottleneck’ will drastically reduce levels of genetic diversity (Meyer & Purugganan, 2013) and was thought to be the norm for cultivated species. However, a relatively long generation time, obligate outcrossing and clonal propagation may mean that hazelnut does not follow this pattern. Furthermore, recent publications have also cast doubt on whether this bottleneck is typical of crops. Emerging evidence suggests that domestication is not a single event but extends over a long period and that the domestication process does not necessarily result in large reductions in genetic diversity (Allaby *et al*., 2019; Smith *et al*., 2019). Given its life history, the large number of cultivars (around 400 clonal cultivars have been described (Thompson et al. 1996)) and smallholdings that maintain them, hazelnut provides a unique opportunity to study the effects of domestication on genetic diversity.

We investigate four main hypotheses surrounding the distribution of genetic diversity in *C. avellana*. We perform clustering analyses and generate summary statistics to test two hypotheses comparing diversity in wild and domesticated hazelnut : (i) There is more genetic structure in cultivated than wild populations and (ii) Domesticated hazelnut have reduced genetic diversity when compared to wild individuals. Before determining how genetic diversity can best be used for crop improvement it must be defined. We sample more than 50 individuals across 17 of the most common cultivars to test whether (iii) Specimens belonging to the same cultivar fall into the same genetic clusters. We then use a variety of approaches to examine test whether (iv) gene flow has occurred between wild and cultivated hazelnut. Finally, we infer phylogenetic relationships among major groups of wild and cultivated hazelnut and estimate the timescale of their divergence to uncover when hazelnut domestication took place.

## MATERIALS AND METHODS

### Sample collection

We sampled putatively wild *Corylus avellana* individuals from 12 sites across Turkey as well as four sites in Georgia and a single site in the UK. Samples of cultivated individuals were taken from locations on the north coast of Turkey and from two sites in southern Italy. A map of collection sites (providing location data were available) in Turkey is shown in Figure 1. Individuals previously identified as *Corylus colurna* and *C. maxima* were sampled from the arboretum at Royal Botanic Gardens, Kew. A full list of samples and their collection locations can be found in Table S1.

**Figure 1.**
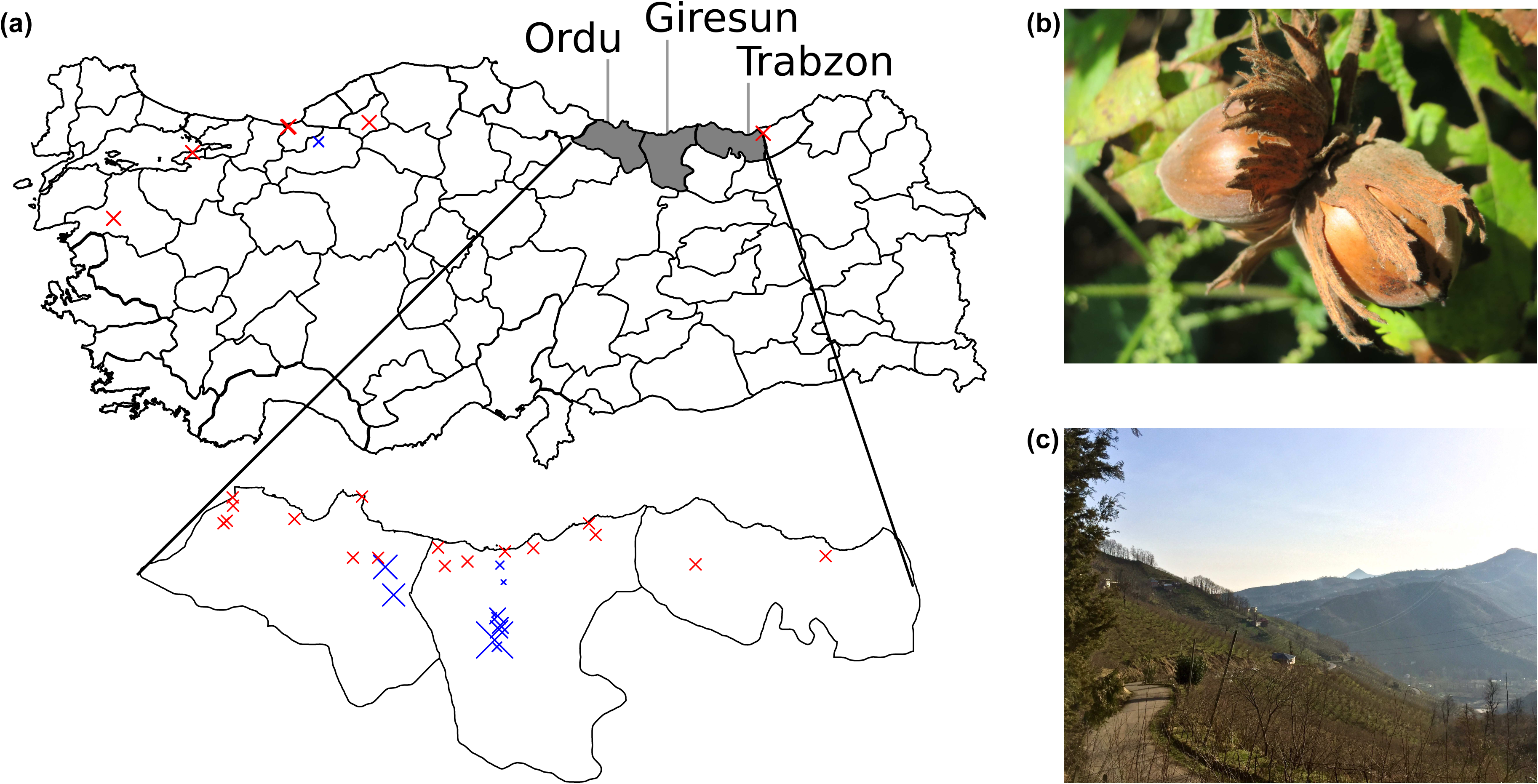
(a) Sampling locations of *Corylus avellana* specimens used in this study. Blue crosses indicate sites where wild individuals were collected and are scaled by number of individuals. Red crosses indicate sites where cultivated individuals were collected, if the information was available. Three major provinces of hazelnut production are highlighted. (b) shows a ripened hazelnut and (c) shows fields of farmed hazelnuts in Giresun. Photo (b) was taken from wikimedia where it was published under a CC0 license and (c) was taken by AJH.

### Library Preparation and sequencing

We extracted Genomic DNA using a modified CTAB mini-extraction protocol (Saghai, 1984; Doyle, 1987). The DNA was then purified using spin columns from the Qiagen DNeasy Plant Mini Kit and then eluted in 60μl water. ddRAD libraries were prepared following Peterson et al. 2012. Briefly, 1 μg of DNA was digested at 37C with the restriction enzyme EcoRI-HF (NEB) for two hours after which MspI (NEB) was added and digestion continued for another two hours. Barcoded adapters (Peterson *et al*., 2012) were ligated to 400 ng digested DNA and samples were pooled. We performed size selection using the Pippin Prep (Sage Biosciences) with a window of 375 to 550bp. We then ran 10 PCR reactions per library to minimize the effect of PCR bias. We repeated this process six times and included two technical replicates each time to check quality across libraries. All libraries were normalised and pooled and then sequenced on four lanes of an Illumina HiSeq 4000 at the Edinburgh Genomics sequencing facility.

### Locus construction and SNP calling

Loci were constructed using STACKS (v1.46) (Catchen *et al*., 2011). We used the program *process_radtags* in to clean and demultiplex reads (options -c -q & -r). Paired-end reads were mapped to a new, draft reference genome for the Turkish cultivar ‘Tombul’ (European Nucleotide Archive (ENA): GCA_901000735) using the Burrows-Wheeler alignment tool (BWA) algorithm (Li & Durbin, 2010) BWA-MEM with the default options keeping only those reads with a mapping quality of 40 or greater. We then used *pstacks* (default parameters) to extract aligned stacks and identify SNPs. We built a catalogue of consensus loci by merging alleles (*cstacks*) based on alignment positions (option -g) and with a maximum of three mismatches allowed between sample loci. We used *sstacks* to search against this catalogue to match loci from each individual to a catalogue locus, again based on alignment position. We then used the *populations* program to filter and output data. We removed loci that were present in less than 75% of individuals and a minor allele frequency threshold of 0.05 was applied; as output, a VCF file was specified to be used for downstream analysis. We then ran a preliminary set of analyses (see below) to detect individuals incorrectly identified as *Corylus*. After this we reran *populations* as above, without misidentified individuals.

### Population diversity and structure

We first performed a principal components analysis (PCA) on the SNP data generated from all individuals and then a discriminant analysis of principal components (DAPC) analysis (Jombart *et al*., 2010) to cluster individuals. The appropriate number of clusters was inferred using Bayesian information criterion (BIC). The number of suitable PCs to retain was identified using the *optim*.*a*.*score* function in ‘adegenet’ (Jombart, 2008).

We then used an alternative clustering approach, fastSTRUCTURE (Raj *et al*., 2014) on our SNP dataset. We ran fastSTRUCTURE with the default settings (which account for admixture) and the simple prior. We used the associated program ‘chooseK.py’ to identify the number of clusters that best explained the structure in the data and the number that maximized the marginal likelihood. We ran analyses using all individuals and then just those identified as domesticated individuals from our DAPC analysis. Results were visualised using the R package ‘pophelper’ (Francis, 2016).

Finally, we ran fineRADSTRUCTURE (Malinsky *et al*., 2018), which uses a different methodology that is based on the fineSTRUCTURE program (Lawson *et al*., 2012). Test runs indicated that including some individuals (e.g. distantly realted *C. colurna* (not including ‘E16’, ‘HAO’ or ‘CK1’) individuals and those with high levels of missing data would yield uninformative results and bias ancestry calculations. These were removed and *popualtions* was rerun, leaving 195 individuals for the final analysis. We filtered our input loci by removing those that had more than 10 SNPs and those that had more than 25% missing data. We ran fineSTRUCTURE with a burn-in of 100,000 steps and then 100,000 further iterations, retaining every 1000^th^.

Summary population genetics statistics were calculated for each cluster inferred using DAPC, fastSTRUCTURE clusters with mixed ancestry individuals removed (to avoid affects of potential admixture) and wild vs. cultivated individuals as differentiated by our fineRADSTRUCTURE analysis. We calculated diversity statistics using functions in the R packages ‘vcfR’ (Knaus & Grünwald, 2016), ‘adegenet’ (Jombart, 2008), ‘hierfstat’ (Goudet, 2005), ‘poppr’ (Kamvar *et al*., 2014) and ‘pegas’ (Paradis, 2010).

### Phylogenetic networks and trees

To understand relationships and distances between samples we used SplitsTree4 (Huson & Bryant, 2005) to infer a phylogenetic network with the neighbour-net algorithm. We used the program PGDSpider (v2.1.1.5; (Lischer & Excoffier, 2012)) to convert the VCF to phylip format, which was used as input. We estimated a network using all samples, include those from *C. colurna* and *C. maxima*.

We also ran SNAPP (Bouckaert *et al*., 2014) to infer a coalescent-based species tree based on binary SNP data. We used the clusters inferred using DAPC as the different taxa. The VCF file was filtered to remove monomorphic loci and only biallelic SNPs were retained. SNAPP is extremely computationally intensive, so to reduce the complexity of our dataset we thinned to SNPs to those with < 3% missing data, used a single SNP per locus and randomly selected five individuals from each of the inferred population clusters. We included *C. colurna* cluster as the outgroup and calibrated the tree using the divergence time between *C. colurna* and *C. avellana* estimated in Helmstetter et al. (Unpublished). A uniform prior was placed on the root where upper and lower bounds encompassed the 2.5/97.5% values of the 95% highest posterior density estimated by Helmstetter et al. (mean = 5.9605, sigma = 0.94). We sampled every 100 generations until convergence (effective sample sizes (ESS) > 200) was reached for all parameters. We assessed convergence using ESS values calculated in TRACER (v1.7; (Rambaut *et al*., 2018)). This process was repeated to ensure that stationarity was reached at the same point across different runs.

### Assessing levels of gene flow among genetic clusters

We used TreeMix to infer patterns of population splitting and mixing from allele frequency data. We calculated allele frequencies for each of the clusters that were identified using DAPC. We sequentially increased the number of migration events from zero to five (m0-m5) and examined changes in likelihood with each event added. We also used the ‘-se’ option to calculate the significance of each migration event. We used two different block sizes (10, 100). We then examined levels of admixture between wild and domesticated clusters using the D statistic (Patterson *et al*., 2012) implemented in the program popstats (Skoglund *et al*., 2015). Significance was calculated using Z scores (D/standard error).

## RESULTS

### Sequencing

On average we recovered 8.21 million retained reads (standard deviation 3.72 million) per sample after processing and cleaning. After identifying and removing incorrectly identified samples our total dataset consisted of 210 individuals. The total SNPs dataset had 64,509 high quality SNPs with an average depth of 79.1 and 13.53% missing data. The large number of SNPs called may be, in part, because we had multiple species in our dataset. All sequences were deposited in the sequence read archive (ENA: PRJEB32239).

### Phylogenetic networks

Our phylogenetic network revealed a clear separation among wild and cultivated individuals (Fig. 2). Generally there was no clear separation among different Turkish cultivars. We were able to identify areas where two major Turkish cultivars, ‘Palaz’ and ‘Tombul’ clustered with other members of the same cultivar. The network revealed a reticulated pattern of branching that linked groups of domesticated individuals, which suggests there is a large amount of conflict in the dataset among cultivars when compared to wild samples.

**Figure 2.**
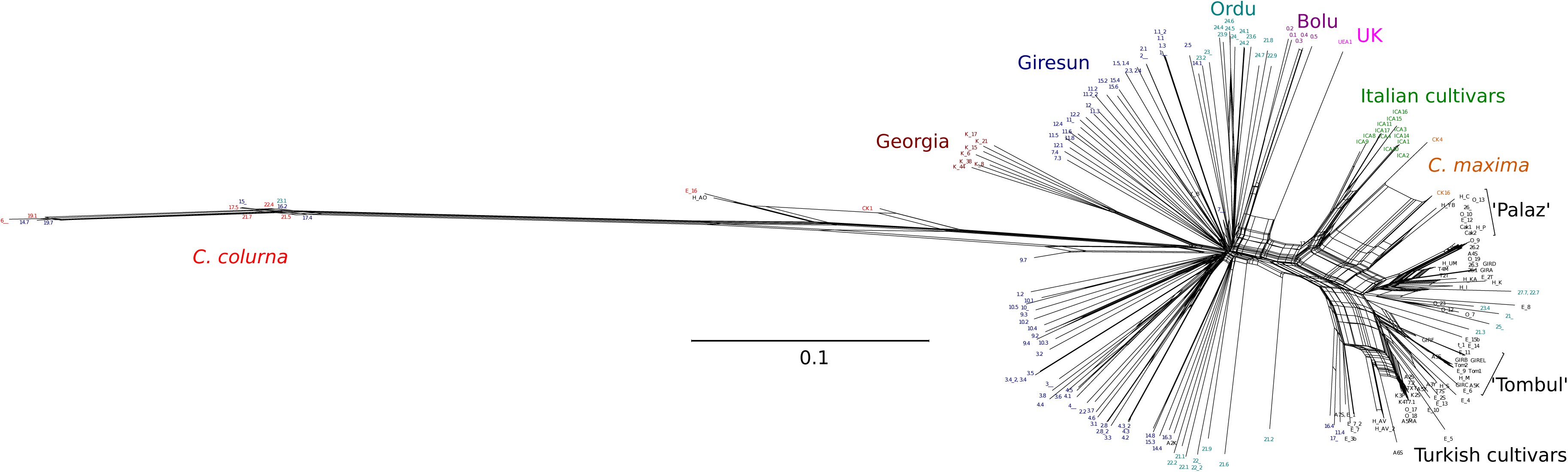
Phylogenetic network calculated using the neighbour-net algorithm across all individuals. A scale is shown inset. Colours at tips correspond to major collection regions or species denoted by group labels of the same colour. Areas where samples from two major Turkish cultivars clustered together are also highlighted.

Distinct groups were more easily distinguishable in wild Turkish individuals. We recovered three major groups corresponding to three different areas of collection, Bolu, Giresun and Ordu (Fig. 2). Samples from Giresun and Ordu were each split into two different groups, indicating that there may be some fine scale genetic structure in these regions. There were a small number of Giresun individuals that fell close to individuals from Ordu, which may point to exchange of DNA between these adjacent regions. Wild Georgian samples were distinct from Turkish individuals, towards the outgroup *C. colurna* while our sole wild individual from the UK was placed in the middle of the split between wild and domesticated samples. Long branches connected *C. colurna* individuals to the major *C. avellana* group. Some individuals originally thought to be *C. avellana* clustered with *C. colurna* and we now consider these as *C. colurna*. Three individuals fell between *C. avellana* and *C. colurna*, one individual considered to be *C. colurna* (E16), a variety of *C. colurna* var. ‘lacera’ and an individual thought to be domesticated *C. avellana* of the cultivar ‘Anac Orta’.

### Population structure

We conducted a DAPC on wild and cultivated individuals together (Fig. 3a) and inferred that six clusters was the optimal number and 13 PCs were retained. Four clusters were made up of cultivated individuals, two of which were markedly different from the others; cluster six contained Italian cultivars (referred to as the Italian cluster) and cluster four contained several individuals of the Turkish cultivar ‘Tombul’ (Turkish cultivars 2, referred to as the ‘Tombul’ cluster). The remaining three clusters were tightly grouped. One of these contained mostly wild *C. avellana* individuals, regardless of their country of origin, Another was made up of Turkish cultivars including many ‘Cakildak’ and ‘Palaz’ (Turkish cultivars 3, referred to as the ‘Cakildak’ cluster). The last cluster of cultivated individuals was a mix of many different strains (Turkish cultivars 1). Although we refer to some clusters by their most prominent cultivar, each also contained a mix of different cultivars. We note that the *C. maxima* samples included in our analysis fell into clusters with cultivated, rather than wild individuals. The final cluster contained individuals previously identified as *C. colurna* as well as those thought to belong to some *C. avellana* cultivars e.g. the cultivar ‘Anac Orta’ (referred to as the *C. colurna* cluster) as in our phylogenetic network (Fig. 2). We treat all members of this cluster as *C. colurna* for downstream analyses. We examined the geographic distribution of the clusters (Fig. 3b) and this revealed evidence for an East-West division between cultivated individuals (‘Cakildak’ cluster and Turkish cultivars 1) along the Black Sea coast.

**Figure 3.**
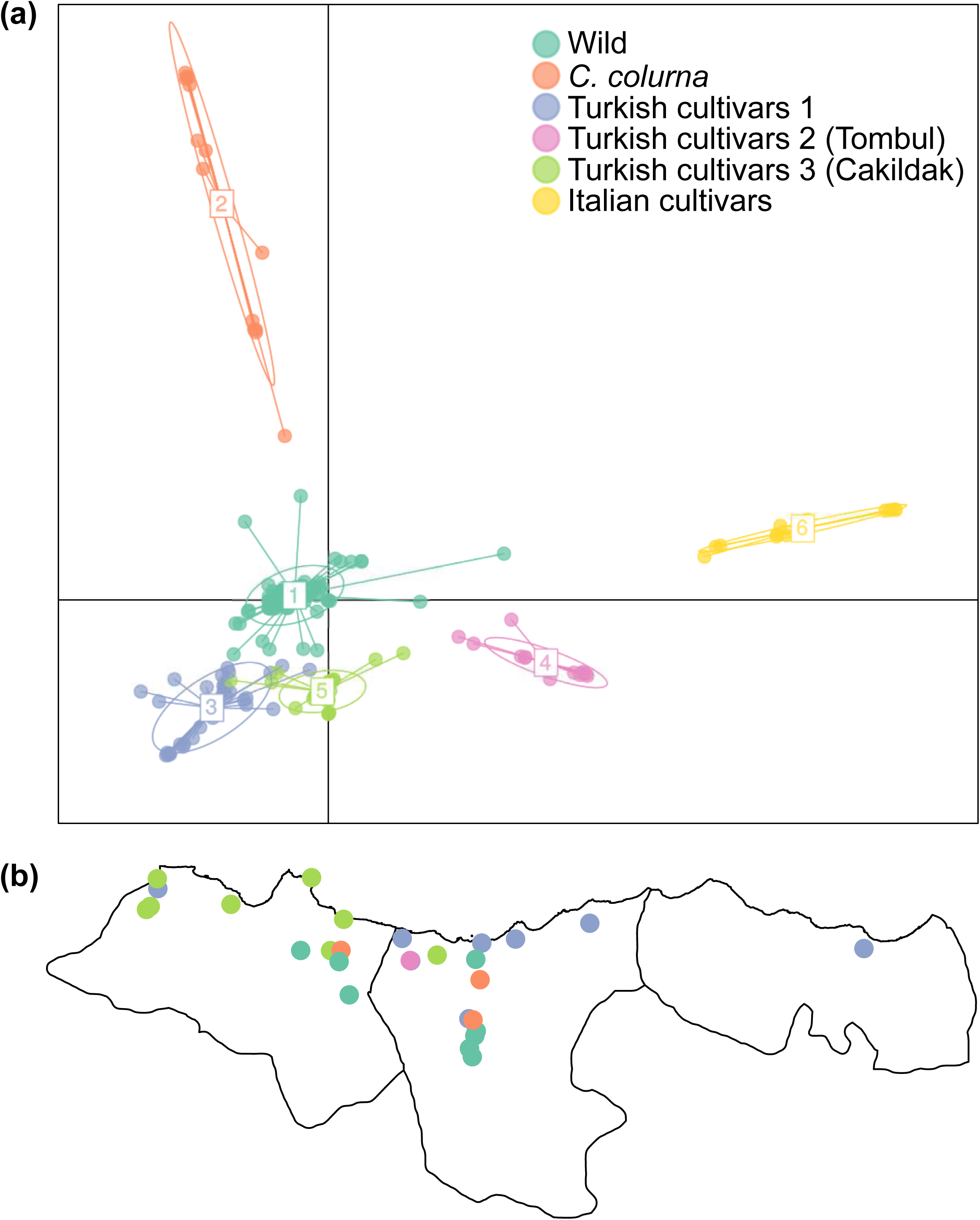
(a) A scatterplot representing showing the locations of wild and cultivated individuals along the first and second axis of our DAPC analysis. The six inferred clusters are labelled and shown in different colours. Cluster 1 primarily corresponds to wild individuals from Turkey, the UK and Georgia. Cluster 2 contains individuals identified as *C. colurna*, Clusters 3-5 contain Turkish cultivated individuals and cluster 6 is made up of Italian cultivated individuals. (b) A map of the Turkish provinces Ordu, Giresun and Trabzon is shown where circles indicate sampling locations (where data was available) and colours correspond to the clusters inferred in (a).

We performed a similar analysis using the same individuals and fastSTRUCTURE. This revealed that eight clusters (k = 8) best explained the structure in the data. Unlike in the DAPC, wild *C. avellana* individuals were spread across multiple clusters. Most fell into a single large cluster (coloured red in Fig. 4c), while groups of individuals from Giresun (teal, Fig. 4c) and samples from Bolu and Giresun (pink, Fig. 4c) also formed distinct clusters of wild individuals. Like in the DAPC analysis, a separate cluster (orange, Fig. 4c) contained individuals identified as *C. colurna* grouped with the same additional *C. avellana* cultivars.

**Figure 4.**
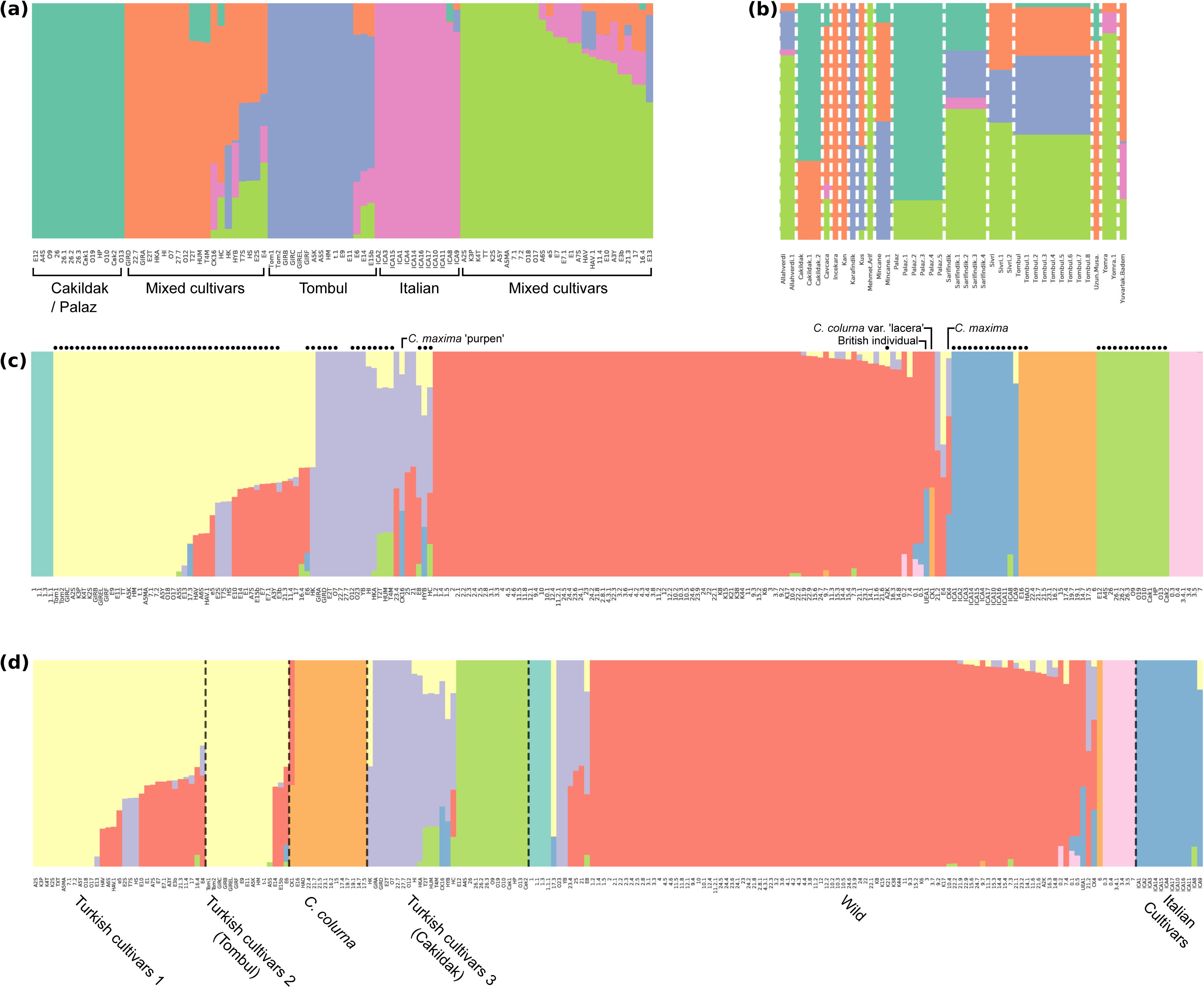
(a) fastSTRUCTURE plot of all cultivated *Corylus avellana* individuals in the dataset. We found that k = 5 best explains structure in the data, which is used in the figure. Major cultivar groups are labelled with the dominant cultivars below the plot. (b) The same analysis as in (a) but individuals with known cultivars are grouped and mean values are calculated for each group. (c) A fastSTRUCTURE plot of all individuals where k = 8 best explained the structure in the data. Black dots indicate those individuals initially identified as domesticated *C. avellana*. Four specific individuals are labelled above the plot. (d) A fastSTRUCTURE plot as in (c) where individuals are grouped based on DAPC clusters (Fig. 3a), as labelled below the plot.

The remaining cultivated individuals were placed into four different clusters. Italian samples grouped together into a distinct cluster. The largest cultivar cluster (yellow, Fig. 4c) in this analysis contained ‘Tombul’ individuals in addition to many other cultivars while the ‘Cakildak’ cluster (green, Fig. 4c) was smaller than in the DAPC analysis. A fourth cluster of domesticated samples (purple, Fig. 4c) again contained a mix of different cultivars. We then grouped our fastSTRUCTURE results using our DAPC clusters (Fig. 4d). This revealed that all fastSTRUCTURE wild clusters belonged to the single DAPC wild cluster. Individuals belonging to Turkish Cultivars 1 and ‘Tombul’ cluster were grouped in fastSTRUCTURE, though most individuals with mixed ancestry were in the former cluster (Fig. 4d). The last major difference between the two analyses was that the ‘Cakildak’ cluster was split in two in the fastSTRUCTURE analysis (Fig. 4d).

The main purpose of this analysis was to uncover evidence of mixed ancestry in wild and domesticated individuals. We detected little evidence for admixture between the *C. colurna* group and other groups, except for the individual ‘CK1’ which was sampled at Royal Botanic Gardens, Kew. This specimen was thought to be a variety of *C. colurna* but may instead be the product of a cross between *C. avellana* and *C. colurna*. We found extensive evidence for admixture among wild and cultivated *C. avellana*. This was particularly evident in two cultivar clusters (yellow and purple, Fig. 4c). We also recovered evidence of admixture between all cultivated clusters, which may be the result of past crosses between cultivars belonging to different clusters. At the same time, there were many domesticated samples with ancestry assigned to just a single genetic cluster, showing little evidence for past admixture.

We also ran a fineRADSTRUCTURE analysis on wild and cultivated individuals. The inferred coancestry matrix (Fig. S1) split wild and cultivated individuals into two separate groups. Many of the wild individuals showed a similar level of coancestry to one another. There were a number of small groups of wild individuals that were grouped by their geographic region – samples from Bolu, Ordu and Georgia shared high levels of coancestry. Individuals from the DAPC *C. colurna* cluster also stood out and were placed within the large group of wild individuals, rather than outside as per expectations. There was a much higher variability in coancestry among cultivated individuals indicating more pronounced genetic structure. They were split into several large groups that broadly reflected the clusters inferred using other approaches, but revealed additional fine-scale structure inside of each group. This approach, alongside others, allowed us to accept our hypothesis that (i) there is more structure in cultivated than wild populations.

### Diversity among wild and cultivated individuals

We found that observed heterozygosity (H_o_) was generally higher in cultivated than wild clusters but estimates of expected heterozygosity (H_e_) did not follow this pattern (Fig. 5). In our assessment of DAPC clusters, wild *C. avellana* had the highest estimated H_e_. This was also true for the largest cluster of wild individuals in our fastSTRUCTURE analysis (Fig. 4c, 5), but the pattern as reversed for the two smaller clusters (Fig. 5). All cultivated clusters had higher H_o_ than wild clusters, across all groups assessed. The ‘Tombul’ DAPC cluster had the lowest H_e_ but in clusters defined by fastSTRUCTURE, one containing ‘Cakildak’ specimens had lower H_e_. When we compared heterozygosity between wild and cultivated individuals as split by fineRADSTRUCTURE (Fig. S1), we found that both H_o_ and H_e_ were similar between the two groups (Fig. 5). Differences between H_o_ and H_e_ indicated that cultivated clusters are typically outbred and wild clusters are inbred. Contrasting patterns of H_e_ and H_o_ meant that we could not accept our hypothesis that (ii) domesticated hazelnut have reduced diversity when compared to wild individuals.

**Figure 5.**
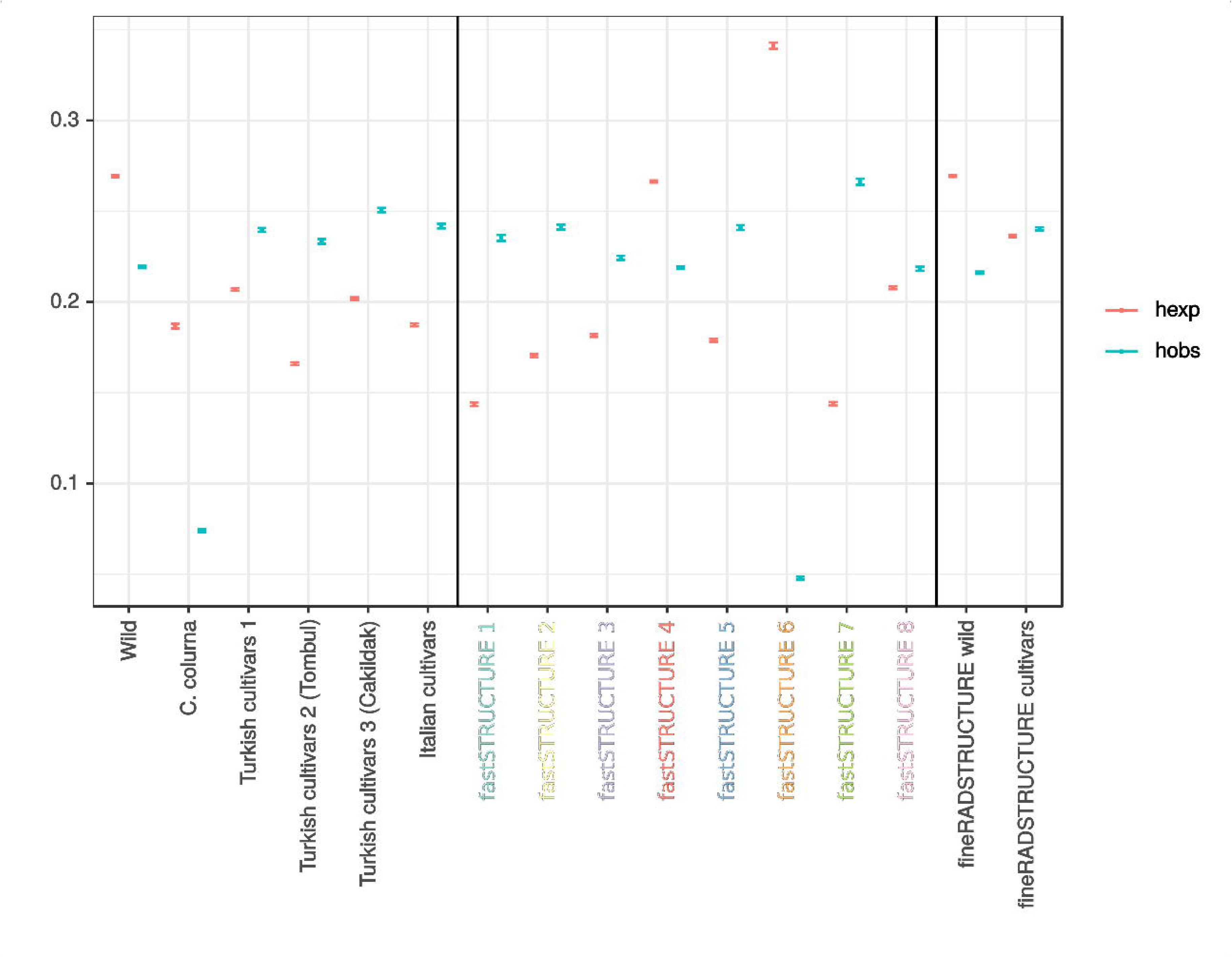
Mean values of expected and observed heterozygosity across all loci (SNPs) showing standard error. We calculated heterozygosity using three different groupings, delineated by black bars. From left to right: the first grouping was based on DAPC clustering (Fig. 3a), the second grouping was based on fastSTRUCTURE clustering and only included individuals with pure ancestry (no admixture) (Fig. 4c). Colours of x-axis labels correspond to the colours used in figure 4c. The third grouping was based on the major split between wild and cultivated individuals in our fineRADSTRUCTURE analysis (Fig. S1).

### Assessing support for predefined cultivars

We aimed to determine whether inferred genetic clusters of cultivated individuals were similar to groups defined by cultivar name. We ran fastSTRUCTURE on cultivated individuals only (‘Tombul’, ‘Cakildak’, Turkish cultivars 1 and Italian clusters from DAPC) and found evidence for extensive genetic structure. Five clusters (Fig. 4a) best explained the structure in the data. These clusters broadly reflected those in the DAPC analyses, except that there were two clusters of mixed cultivars (green and orange, Fig. 4a). Signatures of past admixture between major genetic clusters was inferred in many domesticated individuals, as in the large scale fastSTRUCTURE analysis. Additionally, there was some evidence of admixture involving the cluster of Italian samples, notably in individuals clustered with ‘Tombul’ samples. We then assessed those specimens where the cultivar name information was available by pooling individuals based on cluster name (Fig. 4b). We examined the relative proportion of each cluster that made up each cultivar. For all cases in which we had more than one sample, we found that named cultivars were composed of variation from more than one cluster. We therefore rejected our hypothesis (iii) that genetic clustering supports given cultivar names.

### Phylogenetic relationships and timing of divergence events

After pruning, our final dataset for phylogenetic tree inference consisted of 472 SNPs. Our SNAPP analysis reached convergence (all ESS > 200) after approximately 0.5m generations. The second run converged at the same point after 1m generations, suggesting our results are robust to different starting states. Our SNAPP tree (Fig. 6a) generally had very high support, all but a single node had posterior probability > 0.95. Clusters of Turkish cultivars formed a monophyletic group. The placement of the branch leading to the Italian cultivars was unclear. It was most frequently placed sister to the wild cluster (posterior probability = 0.49; Fig. 6a) but the posterior distribution of trees revealed another relatively common topology in which the Italian cluster was sister to the cluster of wild individuals (Fig. S2), as in our treemix analysis (Fig. 6b). Given our topological uncertainty in the placement of the Italian cluster (Fig. S2), we cannot be certain whether Turkish and Italian hazelnut were domesticated in a single or multiple events. Dating of divergence events indicates that domesticated individuals split from wild individuals between 9.9-16.9kya. The crown age of Turkish cultivars was 5.3-10.2kya and the Italian cluster diverged from wild individuals between 6.5-14.9kya.

**Figure 6.**
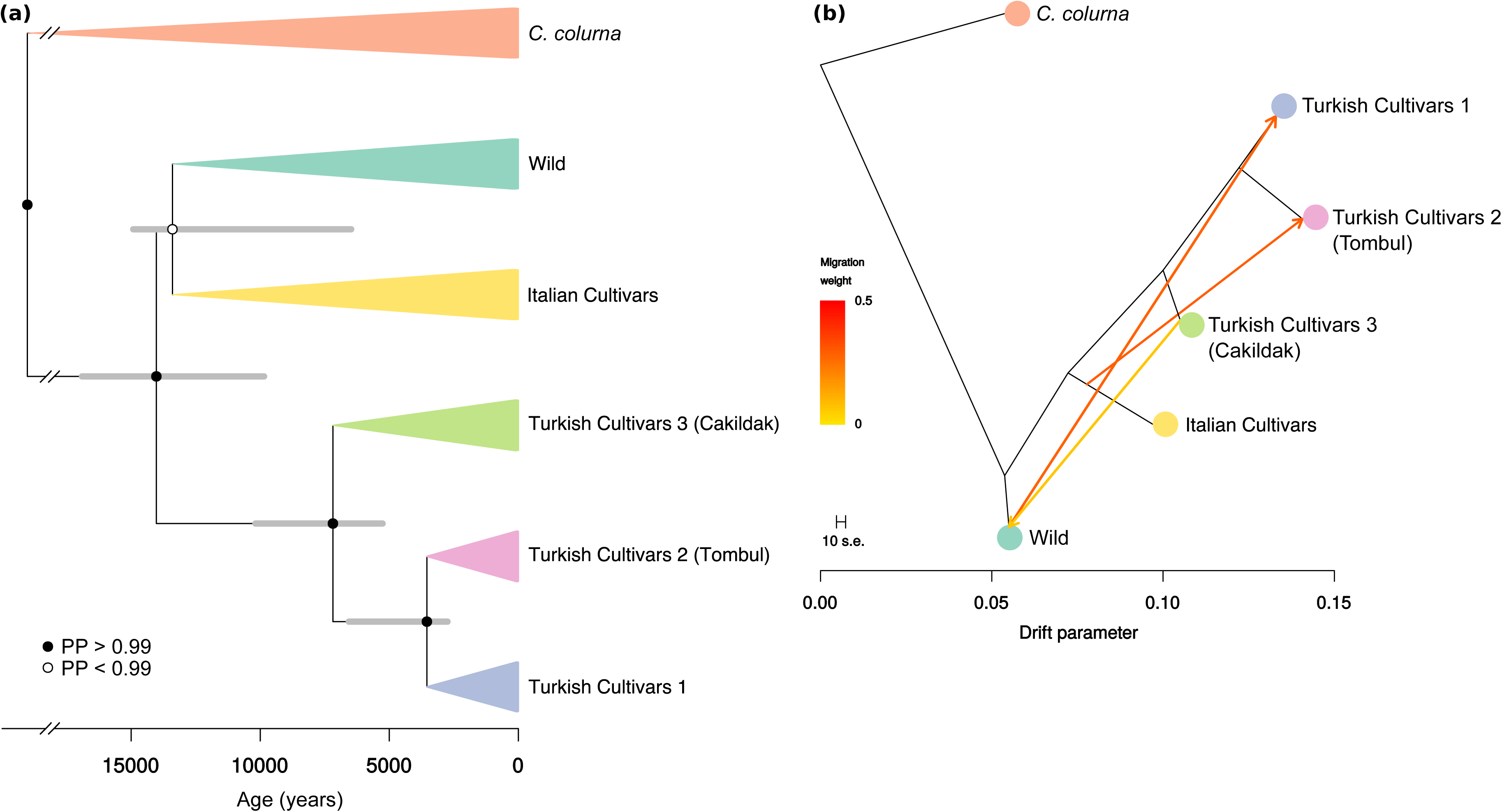
(a) SNAPP tree based on 472 SNPs. Five individuals were randomly selected per DAPC cluster (Fig. 3a). The tree was time-calibrated based on a secondary calibration and an axis is shown below the tree. Inferred 95% Highest posterior densities for node ages are shown as node bars. Branches connected to the root node have been artificially shortened for clarity, so the time axis does not apply beyond the indicated break points. (b) A maximum likelihood tree inferred using TreeMix. The optimal set of three admixture events is also shown on as migration edges, coloured according to their weight, on the tree. Branch lengths are proportional to the amount of drift in allele frequencies among populations, as indicated by the scale. The standard error of the sample covariance matrix is also shown.

### Gene flow among genetic clusters

We used treemix to estimate phylogenetic trees with (Fig. 6b) and without (Fig. S3) migration edges, rooted using the *C. colurna* cluster as an outgroup. The topology of the treemix trees did not place Italian cultivars sister to wild individuals but instead in a clade with the rest of the cultivated clusters (Fig. 6b). We sequentially added migration events, assessing likelihood change at each step (Table. S2) and found that a tree with three migration events had the highest log-likelihood. The first of these migration events went from wild *C. avellana* cluster to Turkish cultivars 1, the second from the Italian cluster to the ‘Tombul’ cluster and third from the ‘Cakildak’ cluster to the wild cluster. The point of origin of a migration event along a branch can indicate whether admixture occurred earlier in time or from a more diverged population, which was the case for the migration event from the Italian cluster. Each of the three events highly was significant (p < 2.1e-06). The amount of variance explained was high (98.24%) even without any migration edges and increased until three migration edges were present, up to 99.98% (Table S2). Matrices of pairwise residuals are shown in Figure S4.

We then examined whether gene flow has occurred between the wild cluster and clusters of Turkish cultivars. We inferred D statistics for three tests (Table S3), two of which had Z scores > 2, indicating some evidence for gene flow between the ‘Cakildak’ and wild clusters, agreeing with our treemix analysis (Fig. 6b). Results from fastSTRUCTURE, treemix and D statistics indicate that gene flow between wild and domesticated hazelnut has taken place and we therefore accept our hypothesis (iv).

## DISCUSSION

### Genetic clusters do not match cultivars

All approaches used revealed that there was more pronounced genetic structure in domesticated than wild hazelnut (Fig. 3, 4, S1). Perhaps the most striking pattern we recovered was the mismatch between genetic data and named cultivars. We identified five genetic clusters across all of our cultivated individuals (Fig. 4a). When we grouped individuals by cultivar name, mean ancestry coefficients were always made up of more than one genetic cluster. This suggests that inferences from our genomic markers do not reflect the naming system of Turkish cultivars. This may be because cultivar names are based on traits that are not correlated with neutral genetic variation, such as kernel size, shape or taste. Morphology has been used to assign Turkish cultivars to three primary groups, primarily based on nut shape (Kafkas *et al*., 2009) and these do not correspond to the genetic clusters we have recovered. Kernels of ‘Yassi Badem’, one of the cultivars that grouped with wild individuals instead of cultivars in our DAPC, are shaped like almonds and not suitable for processing. This cultivar was also found to be the most genetically distant by Kafkas et al. (2009) and did group with cultivars rather than wild individuals in our fastSTRUCTURE analysis (Fig. 4c). It may be that cultivars like ‘Yassi Badem’ have not undergone complete domestication.

Our clustering was similar in some aspects to a previous study based on several nuclear marker types (Kafkas *et al*., 2009). ‘Tombul’ was split among genetic clusters, a pattern also recovered in Boccacci et al. (2006). This cultivar is the most economically important, and it has been implied that it ‘Tombul’ nuts are from just a single clone (Ayfer et al. 1986; Caliskan, 1995) but this is not supported by the genetic variation within ‘Tombul’ we recovered. Furthermore, morphological differences in their nuts and husks have been observed between different ‘Tombul’ samples (Kafkas *et al*., 2009), even while they are still marketed under a single epithet. Kafkas et al. (2009) suggested that Turkish cultivars should be considered as groups of clones with similar phenotypes. Our clustering approach also allows them to be considered by their genetic diversity and shared ancestry. The five clusters of cultivars we inferred provide a helpful starting point for understanding the partitioning of genetic variation across Turkish hazelnut plantations, particularly in light of the potential incompatibilities that could prevent crossing of closely related cultivars. Further work could investigate if any phenotypic traits are associated with these five groups to continue to pave the way for crop improvement.

### Variable distance between domesticated and wild hazelnut

Our DAPC analysis revealed that most cultivated clusters fall close to wild clusters (Fig. 3), an inference that is supported by the work of Ozturk et al. (2017). These patterns could be the result of local domestication, though we think this is unlikely as we would have expected wild and cultivated individuals to cluster together geographically. The ‘Tombul’ and Italian clusters were highly differentiated from other groups in our DAPC (Fig. 3a). Italian cultivars are geographically isolated from Turkish samples as they occur more than 1,500km away, which may explain their differentiation. Boccacci & Botta (Boccacci & Botta, 2009) found little evidence of gene flow from east (Turkey/Iran) to West (Italy/Spain), which supports the differentiation we uncovered. However, we do find some evidence for admixture (Fig. 4, 6b) suggesting that some of the genomes of present day Turkish and Italian cultivars may been the result of past introgression.

The geographic distribution of ‘Tombul’ overlaps with other Turkish cultivars yet it still remains highly differentiated (Fig. 3a), which may be indicative of more considered breeding efforts to improve the cultivar. This cluster also had the lowest level of H_e_ among the six DAPC clusters, suggesting individuals within the cluster are comparatively similar and that this group may consist of only a small number of clones. ‘Tombul’ nuts are considered to be the highest quality so any hybrids may be weeded out by farmers to protect the cultivar. Alternatively, the quality of the nuts may mean that ‘Tombul’ is often planted in new areas where it has not yet had time to interact with local wild relatives. Either way, farmers could be maintaining the distinction between ‘Tombul’ and other cultivars.

### Evidence for gene flow among wild and cultivated samples

We identified two potential instances of past gene flow between wild and domesticated *C. avellana* (Fig. 6b). These were supported by extensive admixture in our clustering analysis (Fig. 4c). However only gene flow between ‘Cakildak’ and wild *C. avellana*, was also supported by D statistic tests. This event was recovered in our treemix analysis (Fig. 6b) and we found some evidence for admixture between wild and ‘Cakildak’ in our fastSTRUCTURE analysis (Fig 4c), which also pointed to extensive admixture between wild *C. avellana* and individuals belong to other cultivars. We also inferred an admixture event between ‘Tombul’ and Italian clusters (Fig. 6c), but was poorly supported by fastSTRUCTURE (Fig. 4a). Overall we have found a complex pattern of recent gene flow between wild and domesticated *C. avellana*.

Crop-to-wild gene flow poses risks relating to the fitness of local wild populations as it can have negative ecological and evolutionary consequences and in some cases even lead to extinction of the wild relative (Ellstrand *et al*., 1999). Conversely, wild-to-crop gene flow may lead to poorer yields if genetic variation underlying traits that have been targeted by breeders is lost. We used a variety of approaches that indicated that introgression -among different cultivars and between wild and domesticated populations - has played a role in generating the diversity we see in domesticated hazelnut in Turkey today. Understanding gene flow between crops and their wild relatives is critical for protecting the local environment and nearby agriculture; our results should prove useful in assessing the impact of these processes in hazelnut.

### A timescale for hazel domestication

Historical documentation of hazel domestication leaves an incomplete picture. As Boccacci & Botta (2009) pointed out, Pliny the Elder (23–79 A.D.) wrote in his work *Naturalis Historia* that the hazelnut came from Asia Minor and Pontus. In the present day, these areas are found on the north coast of Turkey, where our study primarily takes place. The current distribution of *C. avellana* was realised about 7kya, after recolonization following the last glacial maximum (Huntley & Birks, 1983). Between 9-10kya there was a dramatic increase in the amount of pollen found across Europe probably because of nuts dispersed by animals and by human migration. Tribes that existed during the Mesolithic (around 10-6kya) may have been important in the spread of hazel but there is no evidence that they cultivated the plant (Tallantire, 2002).

Our own estimates of the split of cultivated *C. avellana* individuals in Turkey from wild populations (9.9-16.9kya) overlaps with the potential role of early humans in spreading the plant, and may point to propagation. Archaeologists have found an abundance of nutshell fragments during this time period that indicates that hazelnuts were consumed by humans (Bakels 1991; Kubiak-Martens, 1999). It is currently thought that the spread of nuts by Mesolithic humans was by chance (Kuster 2000), but our dating of cultivars splitting from wild populations indicates that this may not have been the case. It is thought that interactions between humans and early crops began in the fertile crescent around 10kya and have continued until the present (Brown *et al*., 2009), similar to our results in hazelnut. Therefore, such an early estimate for the origin of domestication would not be unreasonable and has been found in other crops outside of the fertile crescent (Zheng *et al*., 2016).

Comparisons of sequence data between cultivated and wild individuals can estimate divergence times that predate the origin of the cultivar and are instead closer to the most recent common ancestor for the species (Kim *et al*., 2010; Morrell *et al*., 2011). However, our estimates appear to be too young for a common ancestor of *C. avellana*. Alternatively, changes in generation times through agriculture and strong artificial selection may also change rates of molecular evolution and thus skew divergence times, so our results must be taken with caution. Nevertheless, our estimates suggest that the origin of hazelnut cultivation could predate the Romans and highlights the potential role of Mesolithic tribes in early hazelnut domestication.

### Hazelnut is still in the early stages of domestication

Cultivars are typically expected to have lower levels of genetic diversity (Tanksley & McCouch, 1997) because of the bottlenecks caused by domestication (Eyre-Walker *et al*., 1998) yet we found similar levels of heterozygosity in cultivated compared to wild individuals. This may indicate that the domestication process is still in its early stages, and that any domestication bottleneck has not had a strong effect on genetic diversity. As *C. avellana* is an obligate outcrosser and self-incompatible, any attempts to augment cultivars could also increase levels of heterozygosity. Another possibility is that highly heterozygous individuals have been preferentially retained and clonally propagated in orchards, perhaps because of increased yields caused by hybrid vigour. Our observations are not entirely uncommon: cultivated grapevine (Marrano *et al*., 2017) was more heterozygous than its wild counterpart and a study using microsatellites found that genetic diversity in hazelnut cultivars was similar or higher than wild populations in southern Europe (Boccacci *et al*., 2013).

While levels of H_o_ were lower, levels of H_e_ were actually higher in wild *C. avellana* (Fig. 5), which could point to a reduction of genetic diversity during domestication. We took wild *C. avellana* samples from a wider geographic distribution than cultivated samples and this may have led to the observed patterns of H_e_. Our comparison of all wild and cultivated samples (Fig. S1) accounts for this somewhat, and we find that values of H_o_ and H_e_ are more similar than when using separated clusters (Fig. 5). Furthermore, small clusters of wild individuals inferred using fastSTRUCTURE had levels and patterns of heterozygosity similar to their cultivated counterparts (Fig. 5), so increased H_e_ is not always observed for wild individuals.

Increased heterozygosity is one consequence of introgression and past gene flow between distinct lineages of wild and domesticated *C. avellana* may have contributed to the high levels of H_o_ we observed across cultivars and in turn mask the signal of a domestication bottleneck. However, when we calculated heterozygosity after removing admixed individuals we found very similar results (Fig. 5), which suggests that introgression is likely not driving the observed pattern in genetic diversity. One of the major concerns for modern day crop plants is that reduced genetic diversity caused by domestication will limit the potential for crop improvement in the future (Harlan, 1972). European hazelnut displays relatively high levels of diversity that is promising both for improvement and for resistance to environmental stressors such as pathogens or climate change.

Given the proximity of some wild and domesticated clusters (Fig. 3a), similar levels of heterozygosity (Fig. 5) and existence of cultivars that group with wild individuals, we suggest that hazelnut is still in the early stages of domestication. Our results indicate that cultivated hazelnut may not have experienced a strong domestication bottleneck that reduced genetic diversity. Our phylogenetic analyses suggest that around 10-15kya have passed since domesticated hazelnut first split from its wild progenitors and about 5-10kya since the common ancestor of current Turkish cultivars. This lends support to the idea that domestication has been a gradual process instead of a single event in the past (Brown *et al*., 2009; Brown, 2019), and the genetic proximity of wild and cultivated samples may suggest it is still ongoing today. These characteristics make *C. avellana* a useful model for understanding the genetic effects of partial domestication.

## CONCLUSION

The European hazelnut is one of the most important tree nut crops worldwide and is a large part of the economy and livelihood of communities on the north coast of Turkey. We conducted an assessment of the diversity of cultivars and wild populations in this area and beyond, the first using a genomic approach. We found that cultivars are highly heterozygous, and that admixture has likely occurred among wild and domesticated hazelnut as well as among different genetic clusters of cultivated individuals. We used genomic data to cluster different cultivars into major groups and, surprisingly, these did not overlap with the current naming of cultivars. Our efforts could be useful as a starting point for more efficient use of genetic diversity in breeding programmes. We inferred divergence times of wild and cultivated groups and have estimated a timeframe that aligns with Archaeological evidence for hazelnut consumption in Mesolithic tribes. Our assessment of diversity has provided a new perspective on hazelnut genetics in Turkey and we hope our work will act as a platform for future studies in this economically important crop plant.

## ACKNOWLEDGMENTS

We thank Roberta Gargiulo for the collection of Italian cultivars, Kosta Kereselidze for the collection of Georgian samples and the Hazel Research Centre for providing samples of Turkish cultivars. This work was funded by the British Council’s Newton Fund, grant number: 216394498.

## AUTHOR CONTRIBUTION

RJAB and SJL conceived the study, with input from NO and AJH. SJL, NO and AJH collected samples, NO and AJH conducted molecular lab work. AJH performed data analyses. AJH wrote the initial draft and all authors provided input thereafter.

## SUPPORTING INFORMATION

Additional Supporting Information may be found online in the Supporting Information section at the end of the article.

**Table S1** Collection sites of samples.

**Table S2** Treemix statistics

**Table S3** D statistics

**Fig. S1** fineRADSTRUCTURE coancestry matrix.

**Fig, S2** Posterior distribution of trees from SNAPP analysis.

**Fig. S3** A maximum likelihood tree inferred using TreeMix with no mixture events.

**Fig. S4** Matrices of pairwise residuals from TreeMix analyses.

