## Supplementary Information for "Genetic diversity and domestication of hazelnut (*Corylus avellana*) in Turkey"

Article acceptance date:

The following Supporting Information is available for this article:

**Table S1** Collection sites of samples.

**Table S2** Treemix statistics

**Table S3** D statistics

**Fig. S1** fineRADSTRUCTURE coancestry matrix.

**Fig. S2** Posterior distribution of trees from SNAPP analysis.

**Fig. S3** A maximum likelihood tree inferred using TreeMix with no mixture events.

**Fig. S4** Matrices of pairwise residuals from TreeMix analyses.

**Table S1** Collection sites of samples. Coordinates are given in decimal longitude and latitude for each sample name. In some cases, written descriptions of the location are given.

| sample | latitude | longitude | location description |
| --- | --- | --- | --- |
| UEA1 | 52.621311 | 1.229567 |  |
| ICA15 | 40.807751 | 14.537416 |  |
| ICA17 | 40.807751 | 14.537416 |  |
| ICA10 | 40.814048 | 14.543174 |  |
| ICA11 | 40.814048 | 14.543174 |  |
| ICA16 | 40.814048 | 14.543174 |  |
| ICA4 | 40.814048 | 14.543174 |  |
| ICA9 | 40.888929 | 14.718197 |  |
| ICA3 | 40.885937 | 14.718224 |  |
| ICA2 | 40.884026 | 14.720442 |  |
| ICA1 | 40.890832 | 14.740581 |  |
| ICA8 | 40.897206 | 14.748265 |  |
| ICA14 | 40.899575 | 14.761133 |  |
| 0.1 | 40.808135 | 31.673548 |  |
| 0.2 | 40.808135 | 31.673548 |  |
| 0.3 | 40.808135 | 31.673548 |  |
| 0.4 | 40.808135 | 31.673548 |  |
| 0.5 | 40.808135 | 31.673548 |  |
| O13 | 41.0121 | 37.095183 |  |
| O12 | 41.021633 | 37.111 |  |
| O17 | 41.108483 | 37.139817 |  |
| O19 | 41.108483 | 37.139817 |  |
| O18 | 41.077433 | 37.141883 |  |
| O10 | 41.028167 | 37.443317 |  |
| O23 | 40.883883 | 37.731 |  |
| O9 | 41.111767 | 37.775667 |  |
| O7 | 40.883567 | 37.854183 |  |
| 21 | 40.84899 | 37.889323 |  |
| 21.1 | 40.84899 | 37.889323 |  |
| 21.2 | 40.84899 | 37.889323 |  |
| 21.3 | 40.84899 | 37.889323 |  |
| 21.5 | 40.84899 | 37.889323 |  |
| 21.6 | 40.84899 | 37.889323 |  |
| 21.7 | 40.84899 | 37.889323 |  |
| 21.8 | 40.84899 | 37.889323 |  |
| 21.9 | 40.84899 | 37.889323 |  |
| 22 | 40.84899 | 37.889323 |  |
| 22.1 | 40.84899 | 37.889323 |  |
| 22.2 | 40.84899 | 37.889323 |  |
| 22.4 | 40.84899 | 37.889323 |  |
| 22.7 | 40.84899 | 37.889323 |  |
| 22.9 | 40.84899 | 37.889323 |  |
| 19.1 | 40.884254 | 37.89698 |  |
| 19.7 | 40.884254 | 37.89698 |  |
| 19.7 | 40.884254 | 37.89698 |  |
| 26 | 40.981369 | 37.908017 |  |
| 26.1 | 40.981369 | 37.908017 |  |
| 26.2 | 40.981369 | 37.908017 |  |
| 26.3 | 40.981369 | 37.908017 |  |
| 23 | 40.74486 | 37.930933 |  |
| 23.1 | 40.74486 | 37.930933 |  |
| 23.2 | 40.74486 | 37.930933 |  |
| 23.4 | 40.74486 | 37.930933 |  |
| 23.6 | 40.74486 | 37.930933 |  |
| 23.9 | 40.74486 | 37.930933 |  |
| 24 | 40.74486 | 37.930933 |  |
| 24.1 | 40.74486 | 37.930933 |  |
| 24.2 | 40.74486 | 37.930933 |  |
| 24.4 | 40.74486 | 37.930933 |  |
| 24.5 | 40.74486 | 37.930933 |  |
| 24.6 | 40.74486 | 37.930933 |  |
| 24.7 | 40.74486 | 37.930933 |  |
| 25 | 40.74486 | 37.930933 |  |
| E7 | 40.921391 | 38.149217 |  |
| E11 | 40.852454 | 38.182303 |  |
| E13 | 40.852454 | 38.182303 |  |
| E12 | 40.869386 | 38.293353 |  |
| 16.2 | 40.670197 | 38.424492 |  |
| 16.3 | 40.670197 | 38.424492 |  |
| 16.4 | 40.670197 | 38.424492 |  |
| 2 | 40.577056 | 38.426682 |  |
| 2.1 | 40.577056 | 38.426682 |  |
| 2.2 | 40.577056 | 38.426682 |  |
| 2.3 | 40.577056 | 38.426682 |  |
| 2.4 | 40.577056 | 38.426682 |  |
| 2.5 | 40.577056 | 38.426682 |  |
| 2.8 | 40.577056 | 38.426682 |  |
| 3 | 40.577056 | 38.426682 |  |
| 3.1 | 40.577056 | 38.426682 |  |
| 3.2 | 40.577056 | 38.426682 |  |
| 3.3 | 40.577056 | 38.426682 |  |
| 3.4 | 40.577056 | 38.426682 |  |
| 3.5 | 40.577056 | 38.426682 |  |
| 3.6 | 40.577056 | 38.426682 |  |
| 3.7 | 40.577056 | 38.426682 |  |
| 3.8 | 40.577056 | 38.426682 |  |
| 4 | 40.577056 | 38.426682 |  |
| 4.1 | 40.577056 | 38.426682 |  |
| 4.2 | 40.577056 | 38.426682 |  |
| 4.3 | 40.577056 | 38.426682 |  |
| 4.4 | 40.577056 | 38.426682 |  |
| 4.5 | 40.577056 | 38.426682 |  |
| 4.6 | 40.577056 | 38.426682 |  |
| 1 | 40.551383 | 38.438063 |  |
| 1.1 | 40.551383 | 38.438063 |  |
| 1.2 | 40.551383 | 38.438063 |  |
| 1.3 | 40.551383 | 38.438063 |  |
| 1.4 | 40.551383 | 38.438063 |  |
| 1.5 | 40.551383 | 38.438063 |  |
| 14.1 | 40.666653 | 38.441242 |  |
| 14.4 | 40.666653 | 38.441242 |  |
| 14.7 | 40.666653 | 38.441242 |  |
| 14.8 | 40.666653 | 38.441242 |  |
| 15 | 40.666653 | 38.441242 |  |
| 15.2 | 40.666653 | 38.441242 |  |
| 15.3 | 40.666653 | 38.441242 |  |
| 15.4 | 40.666653 | 38.441242 |  |
| 15.6 | 40.666653 | 38.441242 |  |
| 11.2 | 40.616857 | 38.448466 |  |
| 11.3 | 40.616857 | 38.448466 |  |
| 11.4 | 40.616857 | 38.448466 |  |
| 11.5 | 40.616857 | 38.448466 |  |
| 11.6 | 40.616857 | 38.448466 |  |
| 11.8 | 40.616857 | 38.448466 |  |
| 12 | 40.616857 | 38.448466 |  |
| 12.1 | 40.616857 | 38.448466 |  |
| 12.2 | 40.616857 | 38.448466 |  |
| 12.4 | 40.616857 | 38.448466 |  |
| 6 | 40.856193 | 38.452527 |  |
| 7 | 40.856193 | 38.452527 |  |
| 7.1 | 40.856193 | 38.452527 |  |
| 9.2 | 40.631484 | 38.454621 |  |
| 9.3 | 40.631484 | 38.454621 |  |
| 9.4 | 40.631484 | 38.454621 |  |
| 9.7 | 40.631484 | 38.454621 |  |
| 10 | 40.631484 | 38.454621 |  |
| 10.1 | 40.631484 | 38.454621 |  |
| 10.2 | 40.631484 | 38.454621 |  |
| 10.3 | 40.631484 | 38.454621 |  |
| 10.4 | 40.631484 | 38.454621 |  |
| 10.5 | 40.631484 | 38.454621 |  |
| 17 | 40.792324 | 38.470525 |  |
| 17.3 | 40.792324 | 38.470525 |  |
| 17.4 | 40.792324 | 38.470525 |  |
| 17.5 | 40.792324 | 38.470525 |  |
| E2S | 40.907054 | 38.477369 |  |
| E2T | 40.907054 | 38.477369 |  |
| E3b | 40.907054 | 38.477369 |  |
| E1 | 40.91943 | 38.616417 |  |
| e5 | 41.01171 | 38.889938 |  |
| E10 | 40.968945 | 38.922885 |  |
| E4 | 40.889953 | 40.052355 |  |
| 7.2 | NA | NA |  |
| 7.3 | NA | NA |  |
| 7.4 | NA | NA |  |
| 11 | NA | NA |  |
| 27.7 | NA | NA |  |
| A2K | NA | NA | Cumayani (sea level) |
| A2S | NA | NA | Cumayani (sea level) |
| A3Y | NA | NA | Arabaci koyu (150-200 m from sea level) |
| A4S | NA | NA | Arabaci koyu (150-200 m from sea level) |
| A5K | NA | NA | Aktas koyu Kurtkuyusu mevkii (northern side, 450m from sea level) |
| A5MA | NA | NA | Aktas koyu Kurtkuyusu mevkii (northern side, 450m from sea level) |
| A5S | NA | NA | Aktas koyu Kurtkuyusu mevkii (northern side, 450m from sea level) |
| A5Y | NA | NA | Aktas koyu Kurtkuyusu mevkii (northern side, 450m from sea level) |
| A6S | NA | NA | Aktas koyu Kurtkuyusu mevkii (southern side, 450m from sea level) |
| A7S | NA | NA | Aktas koyu merkez (southern side, 450m from sea level) |
| Cak1 | NA | NA | Yalova |
| Cak2 | NA | NA | Yalova |
| E14 | NA | NA | Not recorded |
| E15b | NA | NA | Not recorded |
| E16 | NA | NA | Giresun, Merkez, GFAE |
| E6 | NA | NA | Not recorded |
| E8 | NA | NA | Not recorded |
| E9 | NA | NA | Not recorded |
| GIRA | NA | NA | Giresun |
| GIRB | NA | NA | Giresun |
| GIRC | NA | NA | Giresun |
| GIRD | NA | NA | Giresun |
| GIREL | NA | NA | Giresun |
| GIRF | NA | NA | Giresun |
| HAO | NA | NA | Not recorded |
| HAV | NA | NA | Not recorded |
| HC | NA | NA | Not recorded |
| HI | NA | NA | Not recorded |
| HK | NA | NA | Not recorded |
| HKA | NA | NA | Not recorded |
| HM | NA | NA | Not recorded |
| HP | NA | NA | Not recorded |
| HS | NA | NA | Not recorded |
| HUM | NA | NA | Not recorded |
| HYB | NA | NA | Not recorded |
| K2S | NA | NA | Not recorded |
| K3P | NA | NA | Yolba_ı |
| K4T | NA | NA | Düzköy-Gelene mah. |
| t-1 | NA | NA | Not recorded |
| T2T | NA | NA | Gençle_tirme bahçesi |
| T4M | NA | NA | Belenköyü-Gö_ceyazı |
| T7S | NA | NA | Den. Bahçesi üstü |
| Tom1 | NA | NA | Yalova |
| Tom2 | NA | NA | Yalova |
| TXT | NA | NA | Not recorded |
| YB | NA | NA | Not recorded |
| CK1 | NA | NA | RBG Kew |
| CK4 | NA | NA | RBG Kew |
| CK16 | NA | NA | RBG Kew |
| K6 | NA | NA | ted 2.3 |
| K8 | NA | NA | ted 3.2 |
| K15 | NA | NA | tsik 1.3 |
| K17 | NA | NA | tsik 2.2 |
| K21 | NA | NA | baz 3.3 |
| K38 | NA | NA | KAKH 1.2 |
| K44 | NA | NA | KAKH 2.2 |

**Table S2** Treemix statistics for different number of mixture events and block sizes.

| No. of mixture events | block size | standard error | % variance explained | ln(likelihood) |
| --- | --- | --- | --- | --- |
| 0 | 10 | 0.00026821 | 0.9824432 | -124.25767 |
| 1 | 10 | 0.00026481 | 0.9913731 | 51.835527 |
| 2 | 10 | 0.00026479 | 0.99921845 | 112.41841 |
| 3 | 10 | 0.00026536 | 0.99980818 | 151.27251 |
| 4 | 10 | 0.00026214 | 0.99976294 | 153.73285 |
| 5 | 10 | 0.0002636 | 0.99964389 | 154.49381 |
| 0 | 100 | 0.0003103 | 0.98079756 | -35.428558 |
| 1 | 100 | 0.00031271 | 0.99046875 | 77.192125 |
| 2 | 100 | 0.0003312 | 0.99760761 | 120.24446 |
| 3 | 100 | 0.00031746 | 0.9998754 | 150.51915 |
| 4 | 100 | 0.00032235 | 0.99964416 | 150.66529 |
| 5 | 100 | 0.0003169 | 0.99985717 | 150.6693 |

**Table S3** D statistics between wild hazelnut and different Turkish cultivars. The D‐statistic compares the occurrence of two site patterns, ABBA and BABA, across a four taxon tree (((P1,P2),P3),O). These patterns can occur by sorting of ancestral polymorphisms, but this would likely result in relatively equal frequencies. If introgression has occurred between P3 and P1 or P2, then one of either ABBA or BABA will occur more often than the other. Significance was calculated using Z scores (D/standard error).

| O | P3 | P2 | P1 | D | Standard errror | z-score | No. sites | No. blocks for jackknife |
| --- | --- | --- | --- | --- | --- | --- | --- | --- |
| colurna | wild_avellana | cult1 | tombul | -0.000432 | 0.00544357 | -0.0793522 | 57548 | 7185 |
| colurna | wild_avellana | tombul | cakildak | 0.01499821 | 0.0060079 | 2.49641311 | 57548 | 7185 |
| colurna | wild_avellana | cult1 | cakildak | 0.0136914 | 0.00514801 | 2.65955443 | 57549 | 7185 |

**Fig. S1** fineRADSTRUCTURE coancestry matrix using all *C. avellana, C. maxima* and those *C. colurna* individuals that were previously identified as domesticated. Darker colours indicate a higher level of coancestry between individuals. Cultivated and wild individuals are clearly separated and there is a higher level of structure amongst cultivated individuals. Clusters of increased coancestry are highlighted on the x-axis.

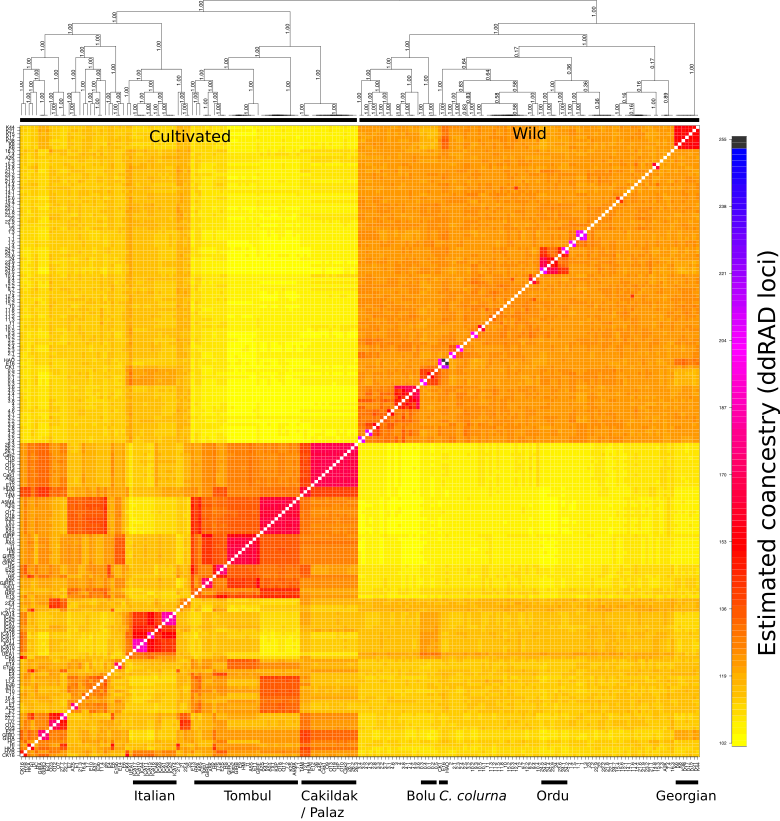

**Fig. S2** Posterior distribution of trees from SNAPP analysis visualized using Densitree. This captures the topological variation encountered during tree inference. The *C. colurna* tip was pruned from all trees. Thin lines represent each tree inferred and thicker lines show the consensus topologies.

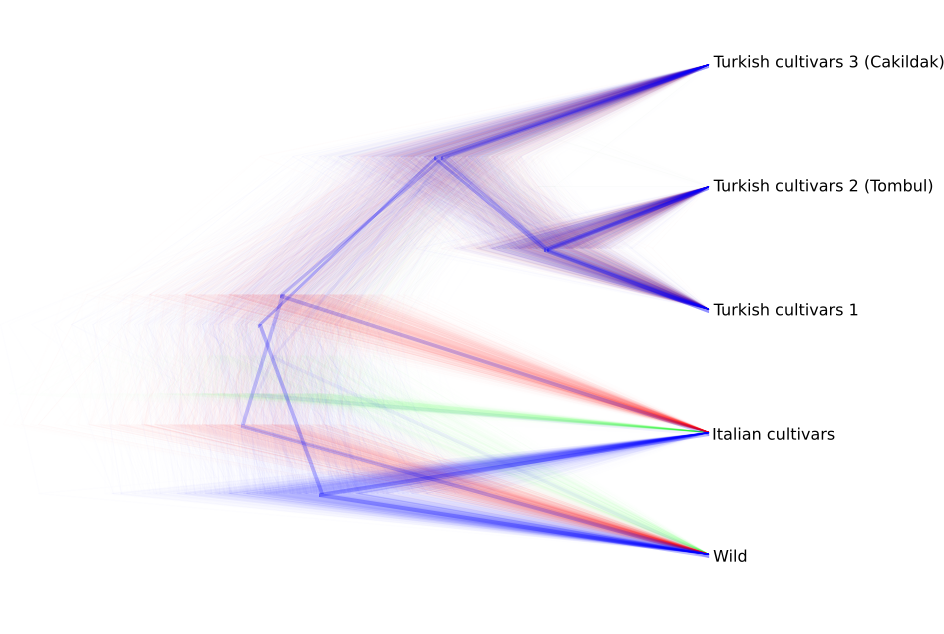

**Fig. S3** A maximum likelihood tree inferred using TreeMix with no mixture events. Branch lengths are proportional to the amount of drift in allele frequencies among populations, as indicated by the scale. The standard error of the sample covariance matrix is also shown.

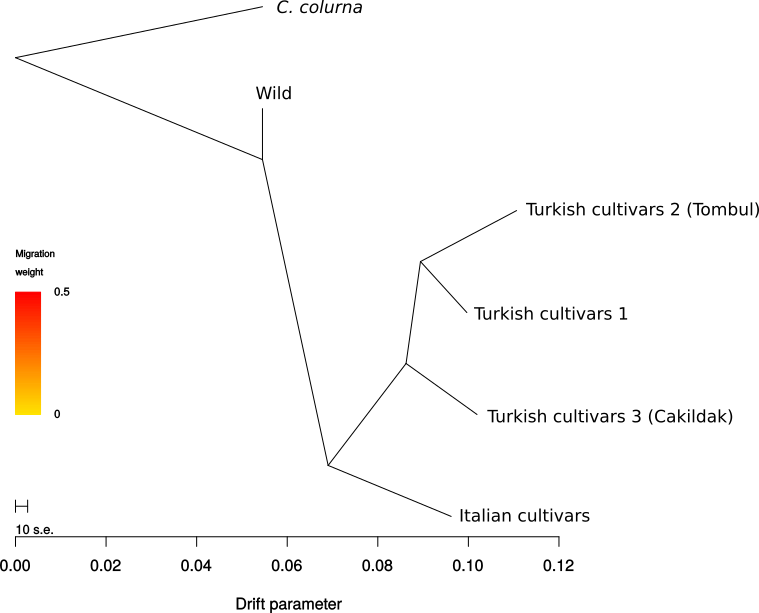

**Fig. S4** Matrices of pairwise residuals for (a) a model with 0 migration edges and (b) a model with 3 migration edges. Larger positive values indicate that there is a poorer fit for the associated population pair.

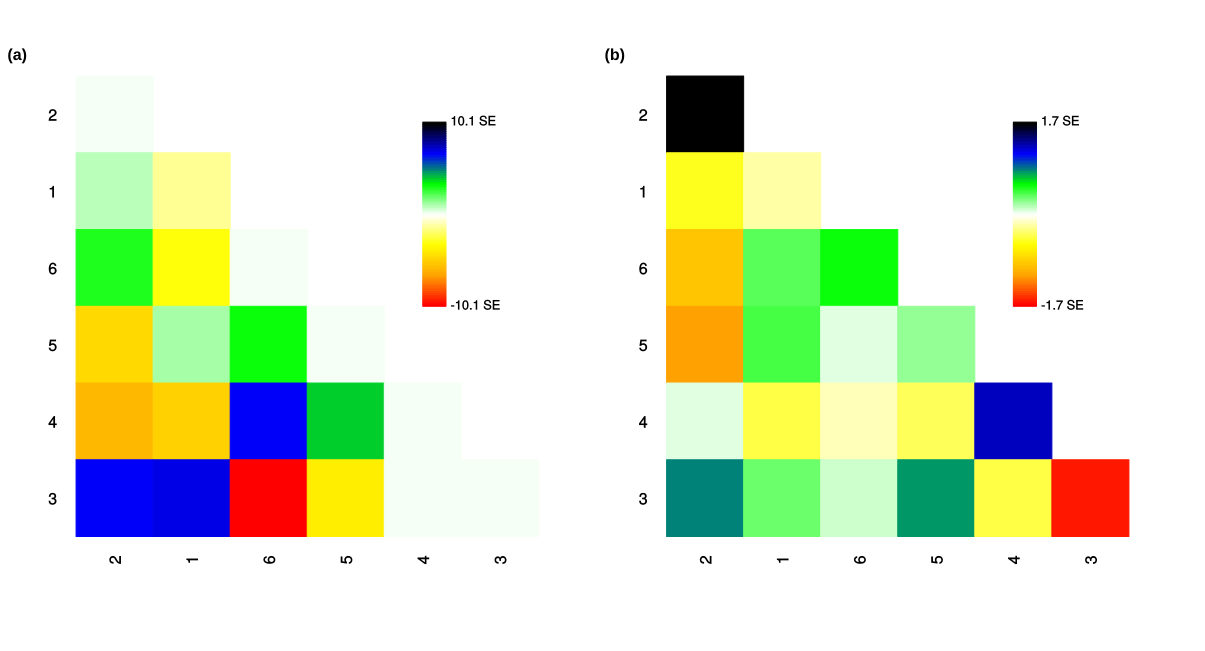
